# Two dominant brain states reflect optimal and suboptimal attention

**DOI:** 10.1101/2020.01.31.928523

**Authors:** Ayumu Yamashita, David Rothlein, Aaron Kucyi, Eve M. Valera, Michael Esterman

## Abstract

Attention is not constant but fluctuates from moment to moment. Previous studies dichotomized these fluctuations into optimal and suboptimal states based on behavioral performance and investigated the difference in brain activity between these states. Although these studies implicitly assume there are two states, this assumption is not guaranteed. Here, we reversed the logic of these previous studies and identified unique states of brain activity during a sustained attention task. We demonstrate a systematic relationship between dynamic brain activity patterns (brain states) and behavioral underpinnings of sustained attention by explaining behavior from two dominantly observed brain states. In four independent datasets, a brain state characterized by default mode network activity was behaviorally optimal and a brain state characterized by dorsal attention network activity was suboptimal. Thus, our study provides compelling evidence for behaviorally optimal and suboptimal attentional states from the sole viewpoint of brain activity. We further demonstrated how these brain states were impacted by motivation, mind wandering, and attention-deficit hyperactivity disorder. Within-subject level modulators (motivation and mind wandering) impacted the optimality of behavior in the suboptimal brain state. In contrast, between-subject level differences (ADHD vs healthy controls) impacted the optimal brain state character, namely its frequency.

## Introduction

Attention is not constant but fluctuates from moment to moment^1-4^. The elucidation of the brain mechanisms required to sustain attention is theoretically important and translationally relevant^5-11^. A wealth of previous studies dichotomized these fluctuations into optimal (stable reaction times, fast reaction times, or fewer attentional lapses/errors) and suboptimal (variable reaction times, slow reaction times, or more attentional lapses/errors) states based on behavioral performance and investigated the differences in brain activity between these behaviorally inferred attentional states^6,7,12-21^. Almost all studies have reported contrasting brain-behavior relationships with task negative networks such as default mode network (DMN) and attention related networks such as dorsal attention network (DAN) and salience network (SN). In a growing number of studies, optimal performance states were associated with greater DMN activity, while suboptimal performance states were associated with greater DAN and SN activity^7,18,21,22^. Although the directionality of the relationship between brain activity and sustained attention is controversial in previous studies, such approaches, which infer mental states based on behavior, are limited by the low dimensionality of behavioral variables—resulting in blunt methods like dichotomization to identify two separate states when in reality, brain dynamics may be more complicated. Furthermore, such approaches rely on frequent responses from participants, and this serves as a strong constraint on the types of tasks that can be used to identify attentional states. To address these stark limitations in the literature, we consider whether attentional states can be defined and observed on the basis of brain activity alone.

Previous studies have also shown relationships between sustained attention ability and the functional connectivity between functionally different brain systems^10,11,18,23^. Functional connectivity is defined as temporal correlation between blood oxygen level dependent (BOLD) time courses from functional magnetic resonance imaging (fMRI). For example, participants who have stronger anticorrelation of functional connectivity between DMN and DAN/SN tend to perform better (more stable and faster reaction times)^19,24^. Individual sustained attention ability also could be predicted by functional connectivity patterns across entire parcellations of the brain (connectome)^10,11,25^. Furthermore, fMRI connectivity neurofeedback training or stimulants such as Methylphenidate have been shown to induce changes in both functional connectivity and sustained attention performance^26,27^. These findings implicate the connectome of functionally different brain systems in supporting sustained attention ability. Nonetheless, whether and how the connectome relates to sustained attention performance through the intermediary of their dynamic brain activity patterns remains unclear^28^.

Here we used a novel energy landscape analysis^29-33^ to identify dynamic brain activity states in a manner that was agnostic to behavior during a sustained attention task. Energy landscape analysis is a data-driven method for estimating stable brain states under the constraints of the connectome between functionally different brain systems. We examined the observed number of states, and whether performance during distinct states corresponded to behaviorally defined optimal and/or suboptimal state(s)^7,18,21^. We provided additional support for these results using an independent validation dataset for replication purposes. To extend this result, we based an additional set of experiments on studies that have shown sustained attention is improved by motivation^34,35^, worsened by mind wandering^22^, and impaired in neuropsychiatric disorders of attention such as attention-deficit hyperactivity disorder (ADHD)^6,8-11^. It remains unknown whether these positive (motivation) and negative (mind wandering, neuropsychiatric disorder) modulators directly impact the composition of the optimal and suboptimal brain states or rather impact the optimality of behavior or likelihood of reaching a given brain state. Using three additional datasets, we investigated how motivation, mind wandering, and ADHD impact the character of these brain states (activity patterns and frequency of these states) and the relationship to performance during these states. Overall, we demonstrate and validate a novel approach for defining attentional states across four independent datasets (see Methods). Further, we provide the first evidence for the existence of and correspondence between neurally and behaviorally defined optimal and suboptimal attentional states. Finally, we demonstrate various influences of positive and negative factors on the character and optimality of these states.

## Results

### Local energy minimum brain states during gradual onset continuous performance task

To investigate the connectome between functionally different brain systems, we first defined 14 region of interests (ROIs) which represent functionally different brain systems (Fig. 1a,b) by applying dictionary learning^36-38^ to resting state fMRI with 16 participants (6 males, ages 18–34 years, mean age = 24.1 years). Dictionary learning can extract brain spatial maps that are naturally sparse and usually cleaner than independent component analysis. We then prepared a time series of average BOLD time courses of 14 ROIs (functionally different brain systems) while 16 participants performed the gradual onset continuous performance task (gradCPT; Dataset 1). GradCPT is a well-validated test of sustained attention, previously used to define attentional states defined by reaction time variability fluctuations over time^7,21^. We then binarized the 14 ROIs’ activity at each time point (by replacing all values above a mean activation with active and others with inactive within each ROI), fitted a pairwise maximum entropy model (MEM)^31,39^ to them, and derived functional connectivity matrices (connectome) between ROIs and average activation of each ROI. Next, based on the connectome between functionally different brain systems and average activation of each ROI, we calculated energy values of all the possible brain activity patterns (2^14^ patterns) (see *Pairwise maximum entropy model* section in the Methods). We then examined hierarchal relationships between the 2^14^ energy values and systematically searched for dominant brain activity patterns that showed locally minimum energy values that were more likely to be observed than similar activity patterns^29,30,32,33^ (Fig. 1c). Please note that this energy value does not indicate any biological energy. It is rather a statistical index that indicates the occurrence probability of each brain activity pattern. For instance, activity patterns with lower energy values tend to occur more frequently. If dynamic brain activity during gradCPT can be described as transitions between behaviorally optimal and suboptimal attention states^21^ (Fig. 1d), such states may correspond to stable brain states which have local energy minimums.

**Figure 1.**
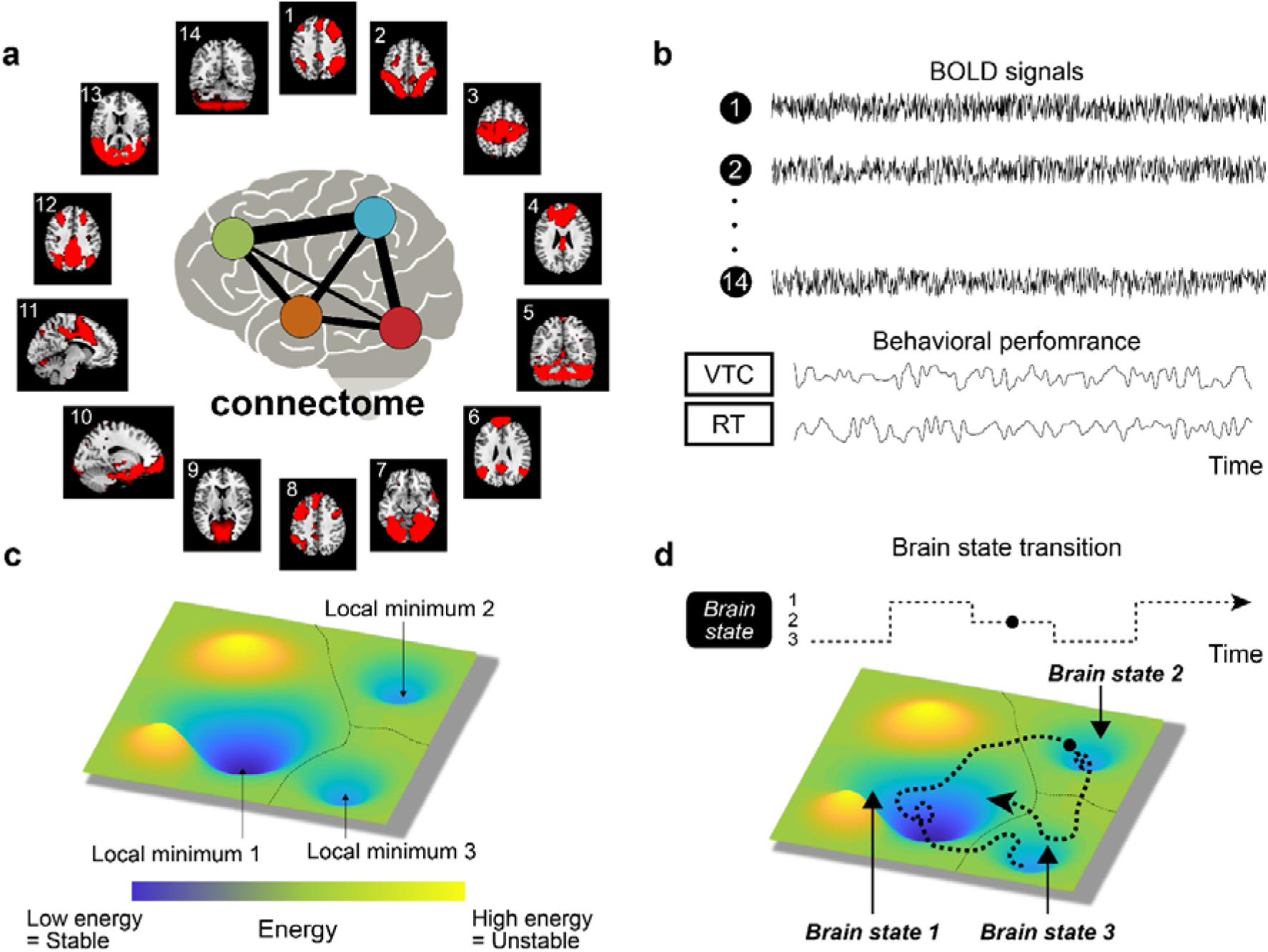
Procedures of energy landscape analysis. (a) Region of interests (ROIs) from dictionary learning using resting-state fMRI. These ROIs indicate functionally different brain regions. Connectome indicates functional connectivity pattern between ROIs. (b) BOLD signals extracted from 14 ROIs and behavioral performances in gradual onset continuous performance task. (c) Energy landscape and local minimums. (d) Brain state transition in energy landscape. BOLD: Blood oxygen level dependent; VTC: Variance time course; RT: Reaction time.

As a result, we found that 13 stable brain states frequently occurred during gradCPT. Figure 2a showed the brain activity patterns in the 14 ROIs for the 13 stable brain states. Figure 2b showed the percentage of dwell time during gradCPT. We found that both State 1 and State 2 occurred about 40% of dwell time during gradCPT indicating that these two brain states were dominant during the task. State 1 was characterized by DMN activity and State 2 was characterized by DAN and SN activity.

**Figure 2.**
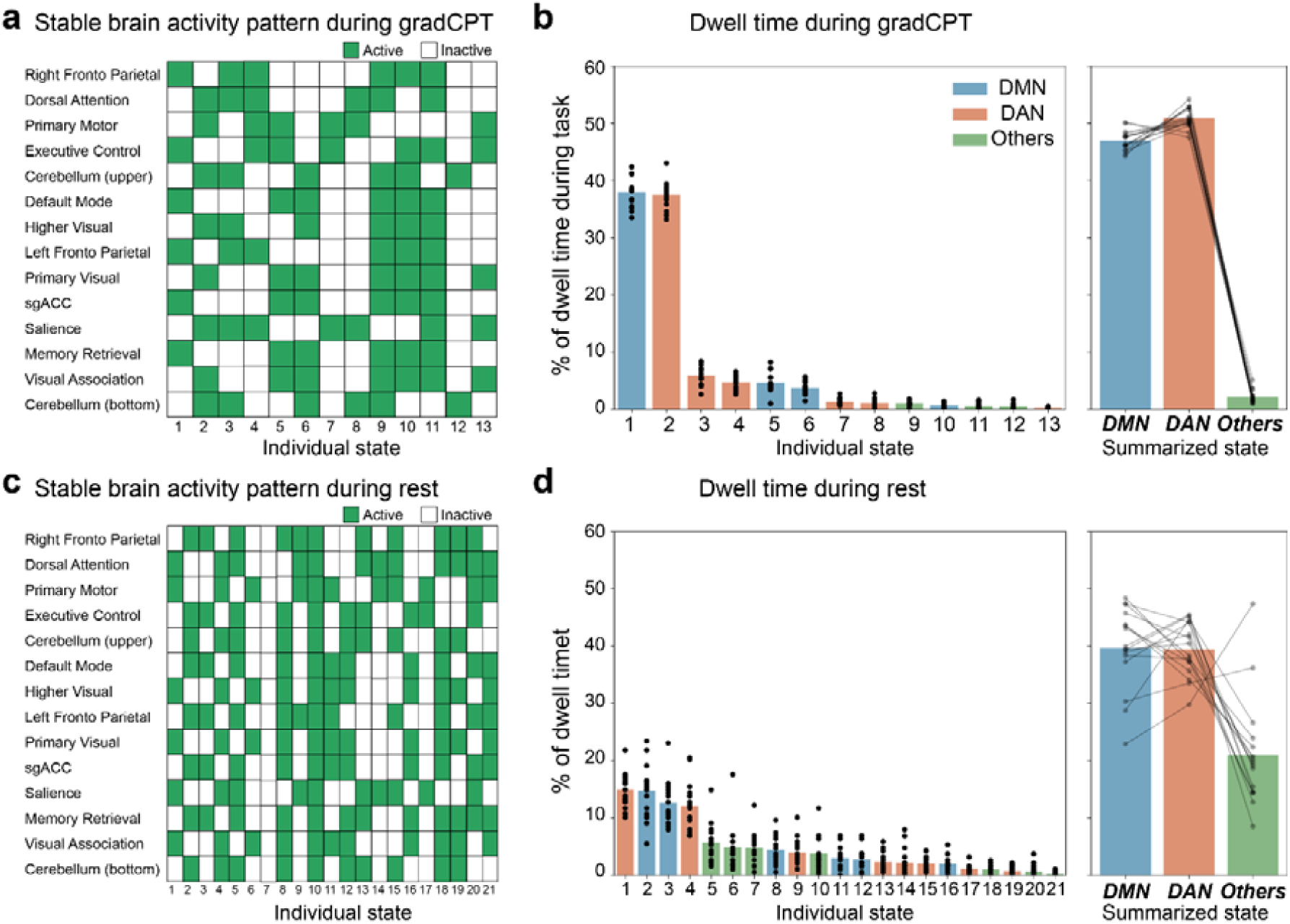
Energetically stable brain states and dwell time during task and rest. (a) Stable brain states during gradual onset continuous performance task (gradCPT). Individual state is represented by an activity pattern in which each brain region is active (green) or inactive (white) state. (b) Left graph shows percentage of dwell time during gradCPT in each individual state. Blue bars show the state defined as DMN-state, red bars show the state defined as DAN-state, green bar show the state defined as Others. Each scatter shows each participant. Right graph shows summation for each summarized state. (c) Stable brain states during resting state. (d) Percentage of dwell time during resting state. sgACC: subgenual anterior cingulate cortex; DMN: default mode network; DAN: dorsal attention network.

We characterized these dominant brain states by focusing on DMN, DAN and SN (ROI 6, 2, and 11, respectively), based on the previous studies linking these functionally different brain systems to fluctuations in sustained attention^7,21^. We then summarized individual local minimum brain states into two major brain state categories (DMN-state and DAN-state) and others. The DMN-state was defined as DMN-active, DAN-inactive, and SN-inactive. The DAN-state was defined as DMN-inactive along with either or both DAN-active and SN-active. If local minimum brain state did not fit into the above criteria, we defined such brain state as “other” (e.g. both DMN and DAN or SN were active or DMN, DAN and SN were all inactive). Using these rules, DMN-state and DAN-state covered 48 % and 51% of total time, respectively (Fig.2b right). We further investigated whether these brain states were specific to the gradCPT. To this end, we investigated local minimum brain states during resting state fMRI. These two brain states existed even in resting state, but these were less dominant, covering 40% and 39% respectively (Figs. 2cd). Furthermore, we confirmed these two brain states were robust to choice of ROIs (Supplementary Figures 2 and 3)^40,41^. For example, when we used the Schaefer et al.^40^ 200 ROIs categorized by networks (7), the only states found were a DMN-state and a DAN+SN-state even without summarizing (Supplementary Figures 2). These results indicate that fluctuation of the brain activity during gradCPT can be described as dynamic transitions between two dominant brain states represented by DMN, DAN and SN activity patterns.

**Figure 3.**
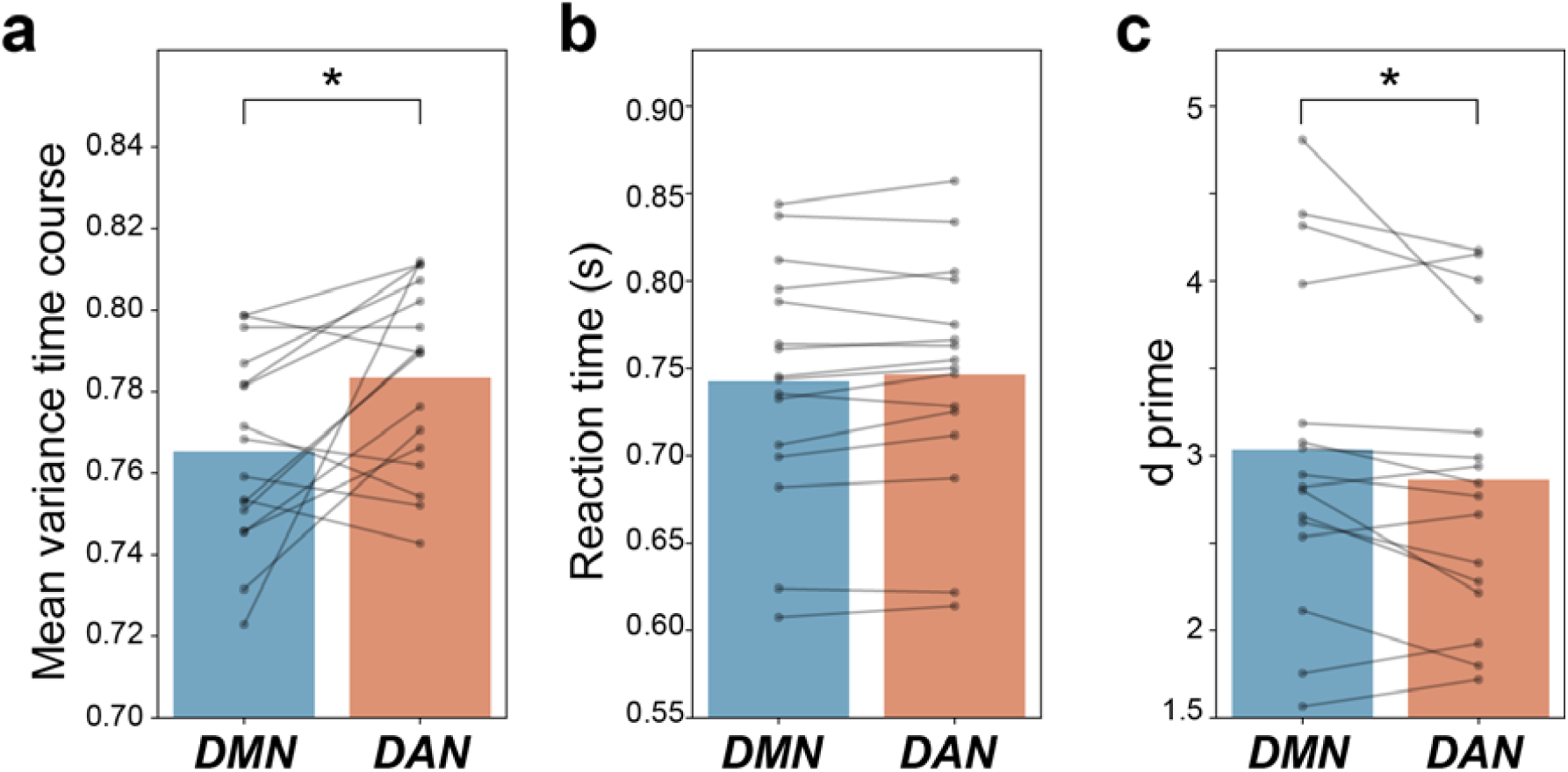
Behavioral performance during each state. (a) Mean variance time course. (b) Reaction time. (c) d prime. Each scatter shows each participant and line connected the same participant. DMN: default mode network; DAN: dorsal attention network. \**P* < 0.05.

We then examined behavioral differences in performances between the DMN-state and DAN-state. We focused on variables commonly used to assess the optimality of sustained attention, including mean reaction time, mean variance time course (VTC, a measure of reaction time variability), and accuracy (d prime) as measures of performance (Fig. 1b). We shifted the time labels of the brain states backwards by 5 seconds to account for the hemodynamic response lag. We found that mean VTC and d prime were significantly better (lower variability and higher accuracy) during timepoints corresponding to the DMN-state than those in DAN-state (Wilcoxon signed-rank test. Variance time course: *W*_*15*_ = 21, *P* < 0.016; Reaction time: *W*_*15*_ = 39, *P* > 0.13; d prime: *W*_*15*_ = 29, *P* < 0.044, two-sided without multiple comparisons) (Fig. 3). Furthermore, we confirmed that the behavioral differences in performances between the DMN-state and DAN-state were robust to choice of ROIs (Supplementary Figures 2 and 3). These results indicate that participants can maintain more stable and accurate performance during DMN-state than those during DAN-state. That is, the DMN-state is a behaviorally optimal state and the DAN-state is behaviorally suboptimal state.

### Replication using an independent validation dataset

We tested whether optimal and suboptimal brain states would generalize to another fMRI dataset in which 29 participants (13 males, ages 21–36 years, mean age = 26.4 years) did five gradCPT runs that had a longer inter-stimulus interval than the original (1300 ms vs. 800 ms per image). Four of five 9-min fMRI runs of gradCPT were gradCPT with thought-probe (see *Long inter stimulus interval (ISI) gradCPT dataset (Dataset 2)* in the Methods section). Here, we did not use the thought-probe measurement (but we did subsequently in *Investigation of the influence of additional cognitive and clinical factors* section). We applied the identical analysis procedure to this independent dataset and found consistent results with the previous experiment that the DMN-state and DAN-state accounted for 49% and 45% of the total scan duration, respectively (Fig.3b right). Although there appears to be 4 main local minimum states (Fig.3a), the pair of brain states 2 and 3 as well as the pair of brain states 1 and 4 are different only in the inclusion or exclusion of the cerebellum ROIs, respectively.

We confirmed that the differences in behavioral performance between during the DMN- and DAN-states largely replicated in this dataset (Wilcoxon signed-rank test. Variance time course: *W*_*28*_ = 57, *P* < 0.00052; Reaction time: *W*_*28*_ = 32, *P* < 0.000061; d prime: *W*_*28*_ = 73, *P* < 0.0018, two-sided without multiple comparisons) (Fig. 3cde). Since this dataset included more subjects and more sessions than study 1 included, or because of the slower ISI, we found that mean RT was also faster in the DMN-state. That is, the DMN-state is a behaviorally optimal state and the DAN-state is behaviorally suboptimal state even in this independent dataset.

### Investigation of the influence of additional cognitive and clinical factors

Next, we investigated how motivation, mind wandering, and ADHD affected sustained attention and the characteristics of these brain states. There are four possibilities for the impact of these factors: (1) the factor directly impact the nature of the brain state(s) (alters the brain activity pattern of brain state), (2) the factor impact the dynamics of the brain state (s) (alters the dwell time in brain state), (3) the factor impacts performance across both brain states equally, (4) the factor impacts performance differentially in one brain state.

#### Influence of motivation

First, we investigated how reward-induced motivation affects performances and brain states using a different fMRI dataset in which 16 participants (10 males, ages = 19–29, mean age = 22 years) performed 3–5 8-min runs of the gradCPT (13 participants completed five runs, 2 completed four runs, and 1 completed three runs) with performance-based rewards (see *gradCPT with reward data set (Dataset 3)* in the Methods). Each 8-min task run was divided into alternating 1-min motivated and unmotivated blocks, which were differentiated by a continuous color border (green for motivated; blue for unmotivated). This yielded 4 min of each block-type per run. During the motivated block, participants earned bonus money for correct responses and lost bonus money for mistakes and during the unmotivated blocks, no money could be gained or lost. These identical reward contingencies were shown to produce reliable improvements in accuracy and RT variability in previous study^34^, thus a priori, we were certain that these performance-based rewards modulated sustained attention performance.

We divided all BOLD signals from 14 ROIs into rewarded and unrewarded blocks and concatenated BOLD signals from all participants for each block. We shifted the time labels of the brain states backwards by 5 seconds to account for the hemodynamic response. We then conducted the energy landscape analysis separately in each block and investigated the stable brain states. As previously found in studies 1 and 2, two dominant brain states were observed for both block types (Supplementary Figure 4). That is, DMN-state and DAN-state were dominant in rewarded blocks (DMN-state: 47%, DAN-state: 52%) and unrewarded block (DMN-state: 49%, DAN-state: 51%) (Fig.5a).

**Figure 4.**
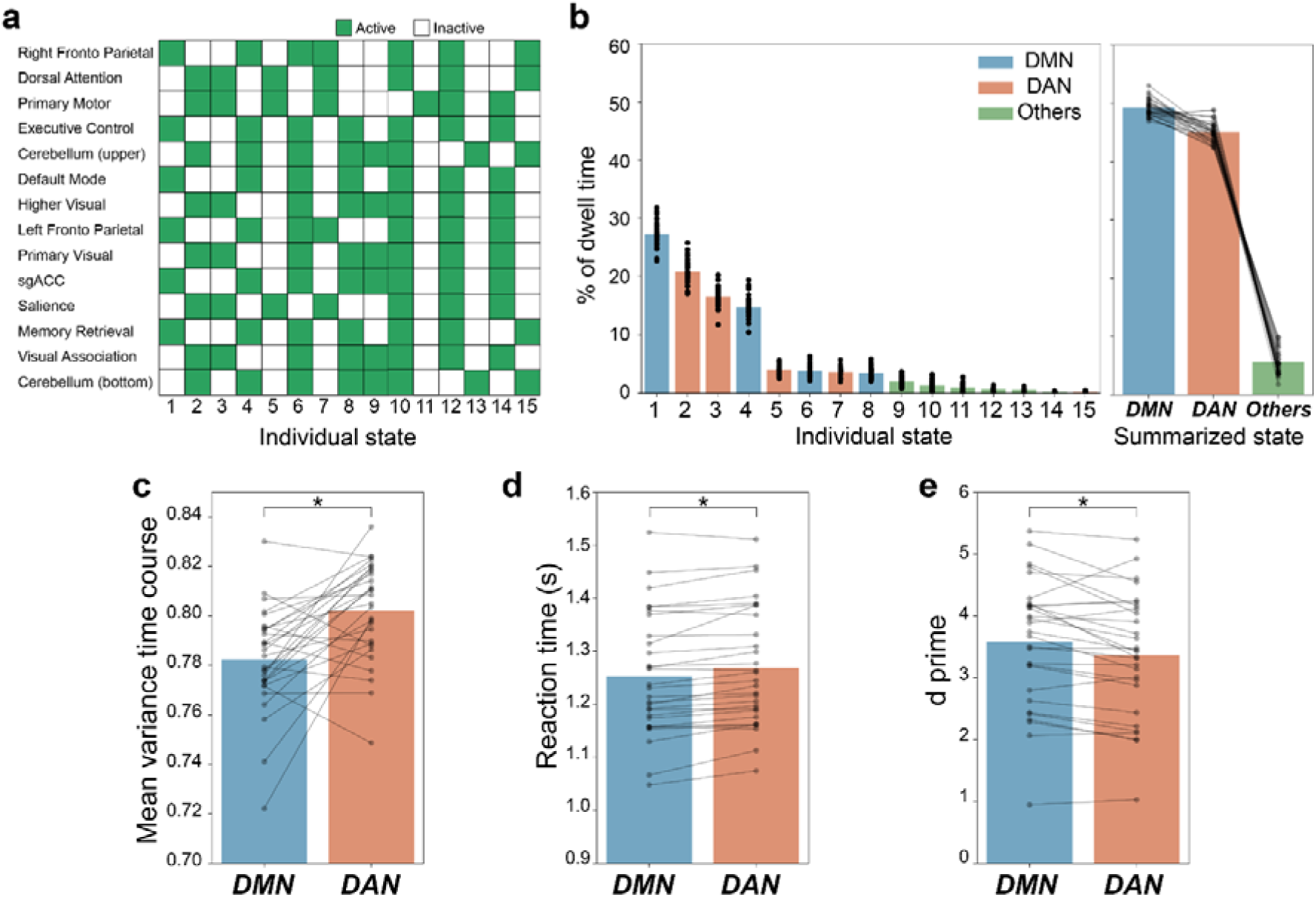
Replication using the independent validation dataset. (a) Stable brain states during long inter-stimulus interval gradual onset continuous performance task (gradCPT). (b) Percentage of dwell time during long inter-stimulus interval gradCPT. (c) Mean variance time course during each state. (d) Reaction time during each state. (e) d prime during each state. Each scatter shows each participant and line connected the same participant. DMN: default mode network; DAN: dorsal attention network. \**P* < 0.05.

We first verified whether the relationship between brain states and behavior could be replicated when using only unmotivated block data, akin to studies 1 and 2 (both without reward). The unmotivated block data is not exactly same as in the first gradCPT dataset because the unmotivated block data might be actively unmotivated by the presence of the motivated block. However, we successfully replicated the difference in performance between DMN-state and DAN-state state (Wilcoxon signed-rank test. mean VTC: *W*_*15*_ = 5, *P* < 0.0012; RT: *W*_*15*_ = 15, *P* < 0.0062; d prime: *W*_*15*_ = 4, *P* < 0.00094, two-sided without multiple comparisons) (Fig.5 pale color). The results using only motivated block data are in the Figure 5 and the legend.

**Figure 5.**
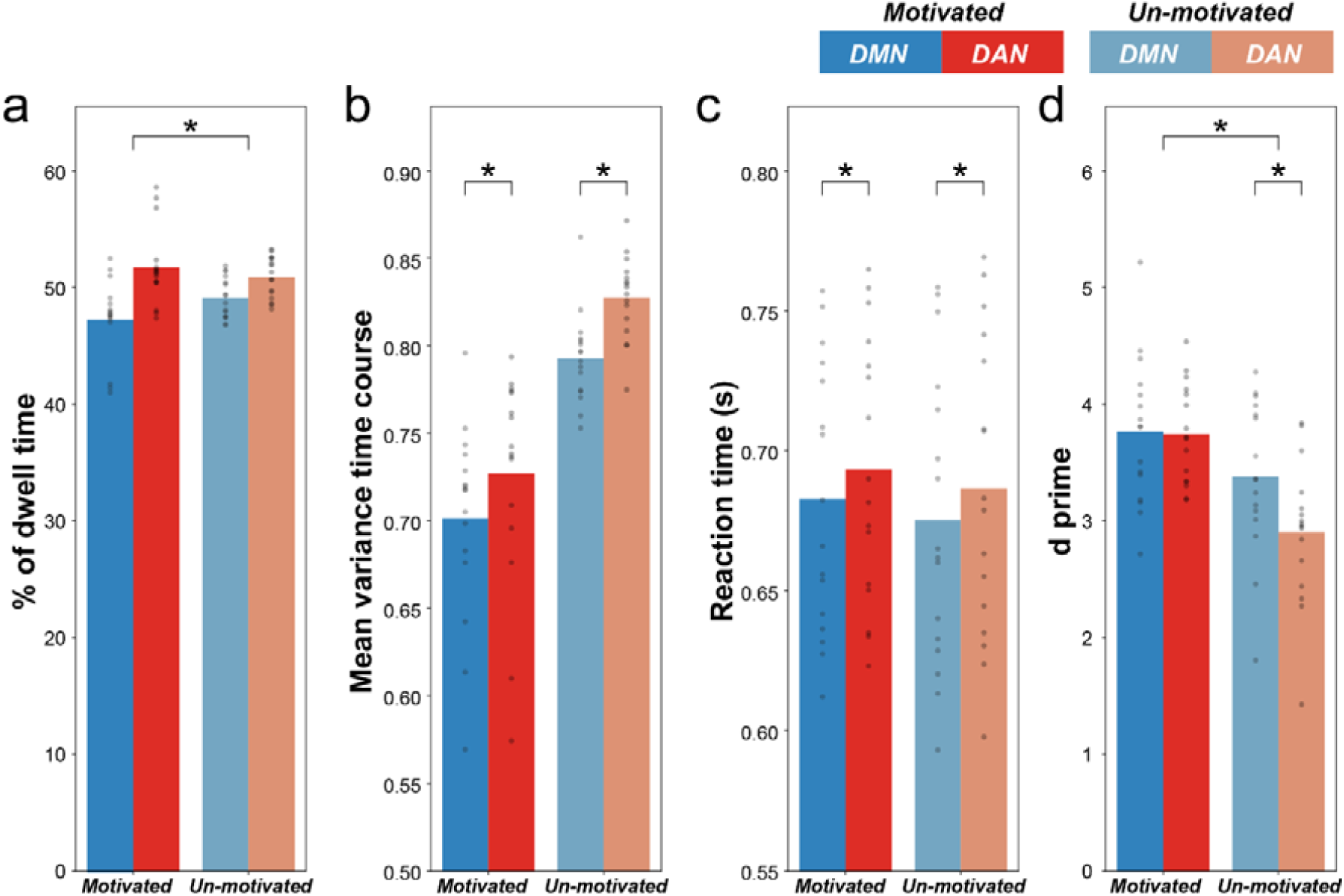
Effect of motivation on dwell time and behavioral performance. (a) Percentage of dwell time during each state. (b) Mean variance time course during each state. (c) Reaction time during each state. (d) d prime during each state. We successfully replicated the difference in performance between DMN-state and DAN-state state in the unmotivated block data and partially replicated in the motivated block data (Wilcoxon signed-rank test. mean VTC: *W*_*15*_ = 24, *P* < 0.03; RT: *W*_*15*_ = 2, *P* < 0.00065; d prime: *W*_*15*_= 53, *P* > 0.43, two-sided without multiple comparison among performances). \**P* < 0.05.

We then investigated whether there were significant interactions between brain state and motivation for dwell time and behavioral performances. We found a significant interaction effect for dwell time as well as d prime (Mixed effects model [interaction effect between brain state and motivation]. Dwell time: *t*_*60*_ = −2.03, *P* < 0.047; d prime: *t*_*60*_ = −2.98, *P* < 0.0042, two-sided without multiple comparisons) (Fig. 5). d prime was significantly better in DMN-state than that in DAN-state during unmotivated block (Mixed effects model [effect of motivation]. *t*_*30*_ = −5.75, *P* < 5.6 × 10^−6^, two-sided after Bonferroni correction for two comparisons [two blocks]), but not during motivated block (Mixed effects model [effect of motivation]. *t*_*30*_ = −0.23, *P* > 0.99, two-sided after Bonferroni correction for two comparisons [two blocks]). This result indicates that motivation could partially overcome the suboptimal brain state’s impact on performance. On the other hand, despite the dwell time interaction, there were no significant dwell time differences between motivated and unmotivated blocks in either brain states (Mixed effects model [effect of motivation]. DMN-state: *t*_*30*_ = 1.91, *P* > 0.12; DAN-state: *t*_*30*_ = −0.95, *P* > 0.68, two-sided after Bonferroni correction for two comparisons [two states]).

#### Influence of mind wandering

Second, we investigated the effect of mind wandering using another fMRI dataset, collected on the same participants from study 2 (part of Dataset 2, see *Long inter stimulus interval (ISI) gradCPT dataset (Dataset 2)* in the Methods). This version of the gradCPT estimated subjects’ self-reported mind wandering degree during gradCPT using experience sampling approach^42^. 29 participants performed 4 gradCPT runs during fMRI, modified here to include thought-probes. Thought-probes appeared pseudorandomly every 44–60 s (three possible block durations of 44, 52, and 60 s). Upon the thought-probe, a question was displayed: “To what degree was your focus just on the task or on something else?” A continuous scale appeared below the question text with far-right and far-left anchors of only task and only else, respectively. Responses were recorded on a graded scale of integers (not visible to the subjects) ranging from 0 (only task) to 100 (only else). This yielded 9 mind wandering degrees every 44-60 s of per run. Following the previous study using this data^22^, we focus on ∼30-s preprobe periods to best yoke thought probe ratings to behaviors. We defined each 30s period as high mind wandering (high MW) periods or low mind wandering (low MW) periods by performing a median split of the 9 mind wandering judgements (high MW period: judgement ≥ median judgement). We concatenated these periods separately and analyzed these as high MW blocks and low MW blocks.

We divided all BOLD signals from 14 ROIs into high MW blocks and low MW blocks and concatenated BOLD signals from all participants for each block. We shifted the time labels of the brain states backwards by 5 seconds to account for the hemodynamic response. We then conducted the energy landscape analysis separately in each block type and investigated stable brain states. Again, we found two dominant brain states for both blocks (Supplementary Figure 5). That is, DMN-state and DAN-state were dominant even in high MW block (DMN-state: 49%, DAN-state: 47%) and low MW block (DMN-state: 49%, DAN-state: 48%) (Fig.6a).

We then investigated whether there were significant interactions between brain state and mind wandering for dwell time as well as behavioral performances. We found significant interactions between brain state and mind wandering for the mean VTC (Mixed effects model [interaction effect between mind wandering and brain state]. *t*_*112*_ = −2.07, *P* < 0.042, two sided without multiple comparisons) (Fig. 6). The difference in the mean VTC between high and low mind wandering level in DAN-state was significantly larger than that in DMN-state (Wilcoxon signed-rank test. *W*_*28*_ = 89, *P* < 0.0055, two-sided). This result indicates that when participants are in the suboptimal brain state, mind wandering particularly impacts performance negatively, namely increasing variability of reaction time.

**Figure 6.**
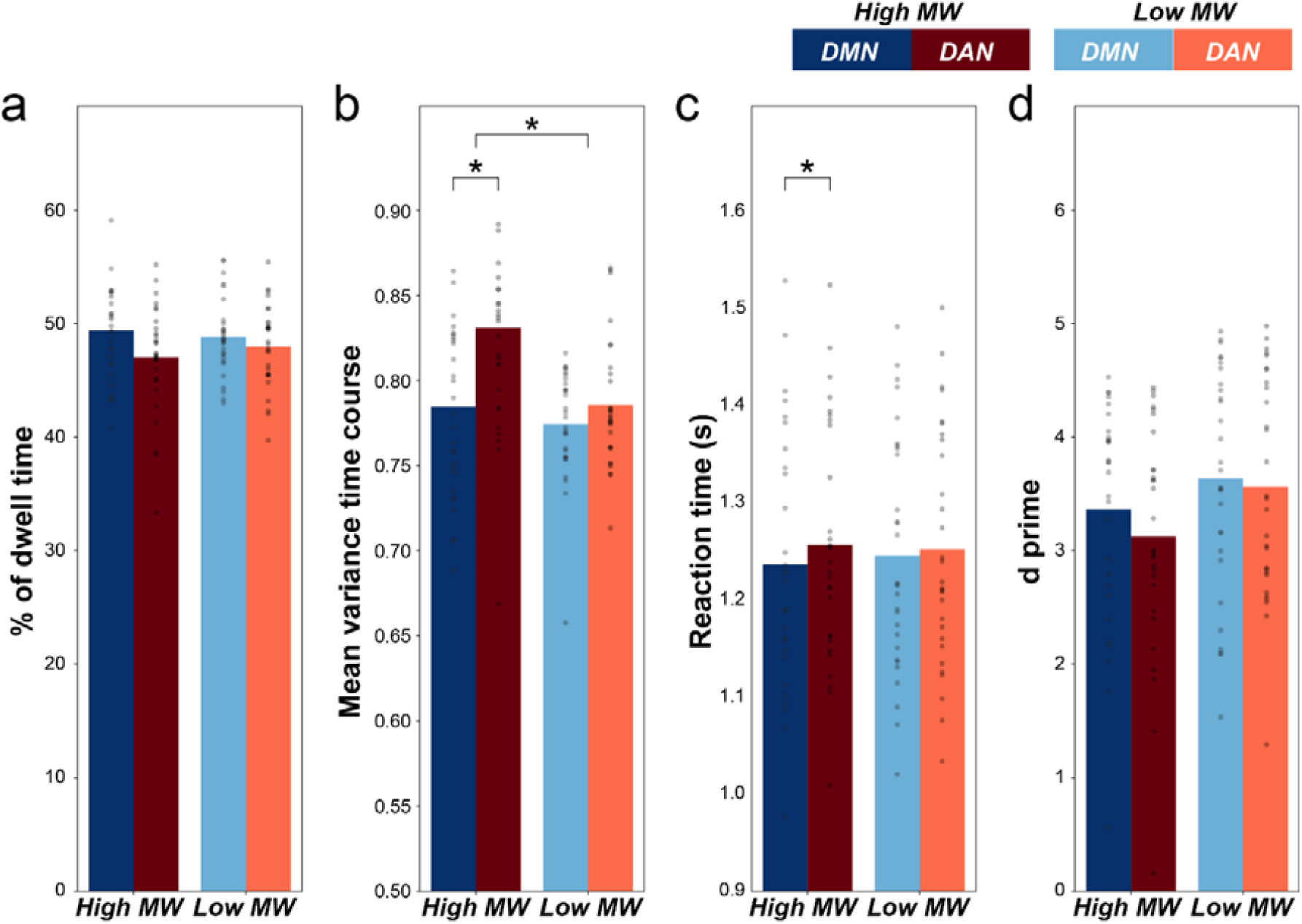
Effect of mind wandering on dwell time and behavioral performance. (a) Percentage of dwell time during each state. (b) Mean variance time course during each state. (c) Reaction time during each state. (d) d prime during each state. \**P* < 0.05.

#### Influence of ADHD

Finally, we investigated the effect of a neuropsychiatric disorder of attention (ADHD) using another fMRI dataset consisting of 19 adult participants with ADHD (8 males, ages 18–34 years, mean age = 24 years) who performed the gradCPT with longer inter-stimulus interval (Dataset 4) (see *Long inter stimulus interval (ISI) gradCPT dataset (Dataset 2)* in the Methods). We applied the identical analysis procedure to this ADHD dataset. We again found two dominant brain states (Supplementary Figure 6). That is, DMN-state and DAN-state were dominant even in ADHD patients (DMN-state: 49%, DAN-state: 47%) (Fig.7a).

We further verified the difference of behavioral performances between during DMN-state and DAN-state (Wilcoxon signed-rank test. Variance time course: *W*_*18*_ = 3.0, *P* < 0.0002; Reaction time: *W*_*18*_ = 10.0, *P* < 0.00063; d prime: *W*_*18*_ = 51.0, *P* > 0.07, two-sided without multiple comparisons) (Fig. 7). Although d prime was not significantly different between DMN-state and DAN-state in ADHD patients, this may be because of lower d prime overall in ADHD patients than that in heathy controls (DMN-state: ADHD d prime = 2.80, HC d prime = 3.57; DAN-state: ADHD d prime = 2.72, HC d prime = 3.36).

**Figure 7.**
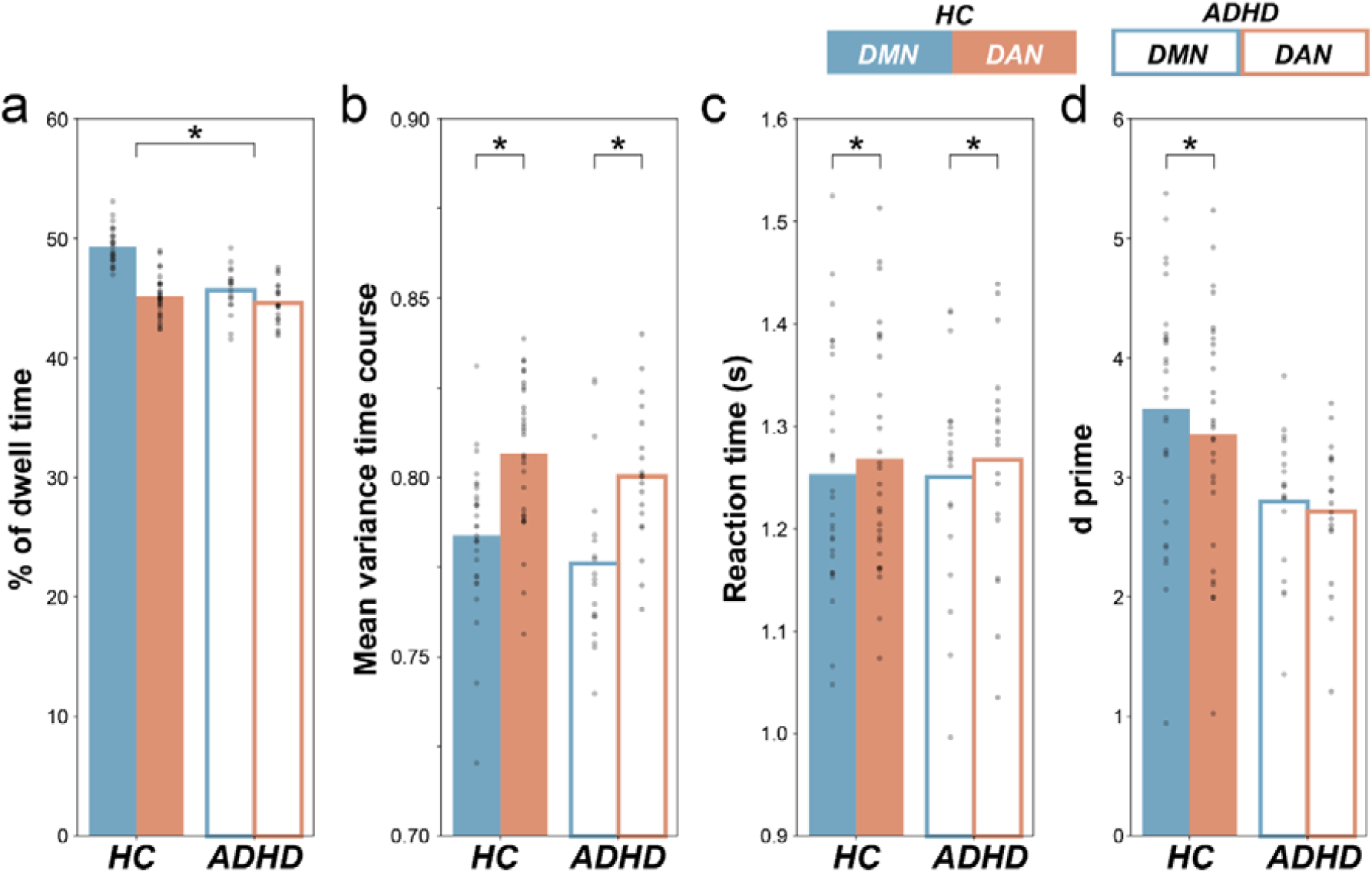
Effect of ADHD on dwell time and behavioral performance. (a) Percentage of dwell time during each state. (b) Mean variance time course during each state. (c) Reaction time during each state. (d) d prime during each state. \**P* < 0.05.

Using study 2 (non-ADHD controls) as a comparison group, we found a significant interaction between brain state and group for the dwell time (Mixed effects model [interaction effect between brain state and group]. *t*_*92*_ = 4.54, *P* < 1.70 × 10^−5^, two-sided without multiple comparisons) (Fig. 7). The dwell times of DAN-state were not significantly different between ADHD and HC (Mixed effects model [effect of group]. *t*_*46*_ = −1.09, *P* > 0.56, two-sided after Bonferroni correction for two comparisons [two states]), however the dwell time of DMN-state was significantly shorter in ADHD patients than in HC (Mixed effects model [effect of group]. *t*_*46*_ = −7.58, *P* < 2.46 × 10^−9^, two-sided after Bonferroni correction for two comparisons [two states]). This result indicates that individuals with ADHD spend less time in the optimal DMN-state than that did HCs, while there was no significant difference in time spent the suboptimal DAN-state.

## Discussion

In the present study, we demonstrated a systematic relationship between dynamic brain activity patterns across the functionally different brain systems and behavioral underpinnings of sustained attention by explaining behavior from observed brain states. This largely confirmed previous findings using behaviorally defined states such that behavioral performance (reaction time variability and accuracy) were superior during a DMN-active state compared to a DAN/SN-active state. That is, DMN-state reflected a more optimal attentional state and the DAN-state reflected a more suboptimal attentional state. We replicated our results in multiple independent datasets. We further revealed the impact of motivation, which does not change the nature and dynamics of these brain-states but serves to overcome performance decrements normally associated with the suboptimal state. Conversely, mind wandering exacerbated the negative impact on performance during the suboptimal state. Finally, we found that individuals with ADHD did alter the dynamics of the brain state by decreasing the dwell time spend in the optimal DMN-state relative to healthy controls.

Our results provided evidence for optimal and suboptimal brain states when defining state independently from behavior. This enabled us to estimate subjects’ attentional states from brain activity without requiring overt responses from subjects. Since it is relatively difficult to get frequent and continuous behavior from many naturalistic tasks, as well as from some patient populations, our attentional states defined by brain activity enhance the ability to track hidden states of attention and could help better reveal neurobiological mechanisms underlying disorders of attention.

In this study, behavioral performance was consistently better during the DMN-state than during DAN-state. This is consistent with several other studies from our lab and others demonstrating DMN activity associated with stable and optimal behavior, and DAN/SN associated with unstable and suboptimal behavior, when states were defined behaviorally^21^. On the other hand, since DMN activity more typically represents “off-task” and is related to mind wandering^43^, it has been thought that DMN activity has a negative impact on performance^17,20^. However, previous studies showed that such relationships are highly complex, such that spontaneous DMN activity can be related to both mind wandering and stable “in the zone” performance^22^. This result indicates that there are multiple possible neural mechanisms for DMN activity. Consistent with this idea, our results did not find significant difference in the time occupied by DMN-state between the high mind wandering blocks and the low mind wandering blocks, suggesting that the difference in the behavioral performance between the DMN-state and DAN-state could be due to a largely distinct mechanisms from mind wandering.

Many previous studies have examined differences in brain activity when behavioral performance was better or worse, without considering the connectome^17,20,21,24^. Studies have also examined the relationship between individual differences in the connectome between functionally different brain systems and sustained attention performance, without considering the brain activity^11,19,23,24^. However, whether and how the connectome relates to sustained attention through the intermediary of their dynamic brain activity has remained unclear^28^. Our energy landscape model clarified this relationship, as it indicates that two stable functional brain systems’ activity patterns with different attentional performance frequently occur under the constraints of connectome.

On the one hand, our results showed that a positive modulator of sustained attention (motivation) partially overcomes the negative effect of the suboptimal DAN-state while a negative modulator of sustained attention (mind wandering) worsens the negative effect in this suboptimal state. This suggest that within-subject fluctuations in performance are more malleable during this suboptimal state. Interestingly, our results showed that when in this suboptimal brain state, motivation improves d-prime while mind wandering increase RT variability. This might suggest that mind wandering and motivation affect cognitive performance during this state via different neural mechanisms (e.g. effect mainly on visual system or on motor system). This is consistent with the interpretation that intrinsic fluctuations of performance are largely distinct from other cognitive mechanisms such as mind wandering and motivation^22,34^. On the other hand, individuals with ADHD spent less time in the optimal brain state than healthy controls, while the relationship between brain states and behavioral performances was comparable across group. These results indicate that within-subject level modulators (motivation and mind wandering) impact the optimality of behavior in the suboptimal brain state, rather than characteristics of the brain state itself. In contrast, between-subject level differences (ADHD vs HC) directly impact the optimal brain state character, namely its frequency. This may indicate that the optimal brain state is less susceptible to positive and negative effects at the intraindividual level, but can be related to interindividual differences in attention ability. As the so-called “in the zone” state may reflect automated information processing^44^ and loss awareness of all other things except for task in progress, the optimal brain state identified in this study may have captured this experience.

The energy landscape created in this study is a type of generative model. Generative models enables us to simulate a transition of brain state when brain network changes (e.g. simulation of drug or connectivity neurofeedback effect) or activity of specific brain regions are inhibited or activated (e.g. simulation of brain stimulation)^26,27,45^. Therefore, in future studies it may be possible to determine optimal targets for neurofeedback and brain stimulation to efficiently remediate and improve sustained attention ability by using our model^27,45-47^.

One limitation of our study is the assumption that brain networks themselves are stable. Recent work suggested a relationship between dynamic functional connectivity and attentional states, thus networks themselves may reconfigure with attentional fluctuations^18,25^.

Novel unsupervised learning techniques, based on Bayesian switching linear dynamical systems (BSDS), provides an integrated framework for identifying latent brain states and dynamic brain connectivity during cognitive tasks^48^. In the future, such advanced techniques could be used to investigate brain states that take into account dynamic functional connectivity and relationship between individual brain state and individual behavioral performance.

In summary, our study is the first to provide evidence for two attentional states, a behaviorally optimal and suboptimal states, from the viewpoint of brain activity. Additionally, our study shows that activity patterns across functionally different brain systems could be the link between prior relationships between functional connectivity (connectome) and sustained attention. Our results indicate that a within-subject level positive modulator (motivation) and a negative modulator (mind wandering) impacted task performance within these brain states, but not the character (composition and frequency) of the brain states themselves. Furthermore, our results suggest that behavior was more susceptible to cognitive modulators of attention when in the suboptimal brain state. On the other hand, our results suggest that while the composition of stable brain states were not different across a between-subject level factor (individual with ADHD vs healthy controls), the time spent in the optimal brain state was shorter in ADHD patients than in healthy controls.

We believe this approach has wide ranging implications for neurocognitive and clinical models of attention, and can set a new methodological and theoretical trajectory for a wealth of future studies.

## Methods

### gradCPT data set (Dataset 1)

#### Participants

Sixteen participants (6 males, ages 18–34 years, mean age = 24.1 years) performed the gradual onset continuous performance task (gradCPT) during functional magnetic resonance imaging (fMRI). The data used in this study and portions of the methods have been published^21^, but the current analyses and results reported have not been published elsewhere. To identify network level functional region of interests (ROIs), a 6-min resting state fMRI was collected and submitted to dictionary learning (details below). All participants were right handed, with normal or corrected-to-normal vision and no reported history of major medical illness, head trauma, neurological, or psychiatric disorder. The study was approved by the VA Boston Healthcare System IRB, and written consent was obtained from all participants.

#### Task Paradigm and Presentation

The gradCPT contained 10 round, grayscale photographs of mountain scenes and 10 of city scenes. These scenes were randomly presented with 10% mountain and 90% city, without allowing the identical scene to repeat on consecutive trials. Scene images gradually transitioned from one to the next, using a linear pixel-by-pixel interpolation, with each transition occurring in 800 ms. Images were projected to participants through a MR compatible goggle system (VisuaStim Digital, Resonance Technology Inc.), and subtended a radius of 2.2° of visual angle. Participants were instructed to press a button for each city scene, and withhold responses to mountain scenes. Response accuracy was emphasized without reference to speed. However, given that the next stimulus would replace the current stimulus in 800 ms, a response deadline was implicit in the task.

#### Behavioral analysis: Reaction time

Reaction times (RT) were calculated relative to the beginning of each image transition, such that an RT of 800 ms indicates a button press at the moment image n was 100% coherent and not mixed with other images. A shorter RT indicates that the current scene was still in the process of transitioning from the previous, and a longer RT indicates that the current scene was in the process of transitioning to the subsequent scene. So, for example, an RT of 720 ms would be at the moment of 90% image n and 10% image n - 1, and so forth. On rare trials with highly deviant RTs (before 70% coherence of image n and after 40% coherence of image n + 1) or multiple button presses, an iterative algorithm maximized correct responses as follows. The algorithm first assigned unambiguous correct responses, leaving few ambiguous button presses (presses before 70% coherence of the current scene and after 40% coherence of the following scene or multiple presses occurred on < 5% of trials). Second, ambiguous presses were assigned to an adjacent trial if 1 of the 2 had no response. If both adjacent trials had no response, the press was assigned to the closest trial, unless one was a no-go target, in which case subjects were given the benefit of the doubt that they correctly omitted. Finally, if there were multiple presses that could be assigned to any 1 trial, the fastest response was selected. Slight variations to this algorithm yielded highly similar results, as most button presses showed a 1–1 correspondence with presented images.

#### Behavioral analysis: Variance time course

Beyond mean RT and error rates, we were particularly interested in trial-to-trial variation in RT, which we assessed via a novel within subject analysis that we called the variance time course (VTC)^21^. VTCs were computed from the ∼500 correct responses in each run (following z-transformation of RTs within-subject to normalize the scale of the VTC), where the value assigned to each trial represented the absolute deviation of the trial’s RT from the mean RT of the run. We reasoned that deviant RTs, whether fast or slow, represented reduced attention to the task as follows: extremely fast RTs often indicate premature responding and inattention to the potential need for response inhibition ^49^, while extremely slow RTs might indicate reduced attention to or inefficient processing of the ongoing stream of visual stimuli, requiring more time to accurately discriminate scenes^17^. Values for trials without responses (omission errors and correct trials) were interpolated linearly, such that the missing values were linearly estimated from RTs of the 2 surrounding trials. A smoothed VTC was computed using a Gaussian kernel of 9 trials (∼7 s) full-width at half-maximum (FWHM), thus integrating information from the surrounding 20 trials, or 16 s, via a weighted average. This choice was based on prior work linking fluctuations around this frequency to attentional impairments^50^.

#### MRI Acquisition

Scanning was performed on a 3T Siemens MAGNETOM Trio system equipped with a 12-channel head coil, at the VA Boston Neuroimaging Research Center. Functional runs included 248 (gradCPT) or 188 (resting state) whole-brain volumes acquired using an echo-planar imaging sequence with the following parameters: repetition time (TR) = 2000 ms, echo time (TE) = 30 ms, flip angle = 90°, acquisition matrix = 64 × 64, in-plane resolution = 3.0 mm^2^, 33 oblique slices, slice thickness = 3, 0.75 mm gap. MPRAGE parameters were as follows: TE = 3.32, TR = 2530 ms, flip angle = 7°, acquisition matrix = 256 × 256, in-plane resolution = 1.0 mm^2^, 176 sagittal slices, slice thickness = 1.0 mm.

#### fMRI analysis: Preprocessing of fMRI

We performed preprocessing of the fMRI data using FMRIPREP version 1.3.0 ^51^. Preprocessing steps included realignment, coregistration, segmentation of T1-weighted structural images, normalization to Montreal Neurological Institute (MNI) space. For more details of the pipeline, see http://fmriprep.readthedocs.io/en/latest/workflows.html.

#### Parcellation of brain regions: Dictionary learning

We identified functionally different brain systems to use as functional ROIs from the whole brain by applying dictionary learning to resting state fMRI data ^36-38^. We first concatenated all participants’ resting state fMRI and then applied dictionary learning implemented in Nilearn^52^. Dictionary learning is a sparse based decomposition method for extracting spatial maps. We set the number of components as 20. We used spatial smoothing with an isotropic Gaussian kernel of 6 mm full-width at half-maximum. A temporal bandpass filter was applied to the time series using a first-order Butterworth filter with a pass band between 0.01 Hz and 0.08 Hz to restrict the analysis to low-frequency fluctuations, which are characteristic of resting state fMRI BOLD activity^53^. The BOLD signal time courses were extracted from these 20 ROIs. We visually inspected 5 ROIs which were considered as noise and an auditory related ROI which were not related to our current task and excluded from our current analysis. We finally used 14 ROIs for all analysis (Fig. 1a).

#### Physiological noise regression

Physiological noise regressors were extracted by applying CompCor^54^. Principal components were estimated for the anatomical CompCor (aCompCor). A mask to exclude signals with a cortical origin was obtained by eroding the brain mask and ensuring that it contained subcortical structures only. Six aCompCor components were calculated within the intersection of the subcortical mask and union of the CSF and WM masks calculated in T1-weighted image space after their projection to the native space of functional images in each session. Furthermore, to isolate the effect of each trial type (commission error, correct omission, correct commission, omission error) as well as trial-to-trial RT, we included mean evoked response for each trial type and trial-to-trial RT. We estimated BOLD response time courses of each event type by using *hemodynamic_models* function implemented in Nistat (https://nistats.github.io/). To remove several sources of spurious variance, we used linear regression with eighteen regression parameters, including six motion parameters, average signal over the whole brain, six aCompCor components, estimated BOLD response time course of each event type, and estimated BOLD response time course of trial-to-trial RT.

#### Pairwise maximum entropy model

We fitted the pairwise Maximum entropy model (MEM) to the preprocessed BOLD signals as follows in the same manner as that employed in the previous studies^29-33,39^. We used open toolbox so called Energy Landscape Analysis Toolkit (ELAT) (https://sites.google.com/site/ezakitakahiro/software). Since this method cannot be applied to relatively large amount of data^30^, to achieve a high accuracy of fitting we had to reduce the number of ROIs to 14 ROIs of 20 ROIs excluding 6 ROIs which seemed to be visually noise or auditory related components. For each ROI, we first binarized the obtained fMRI signals with a threshold that was defined as the time-averaged activity of the same ROI. We then concatenated BOLD signals from all participants for each ROI. Previous studies suggest that binarization does not eliminate important information contained in originally continuous brain signals ^31,32,39^. In this method, the binarized activity 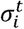 at ROI *i* and discrete time *t* is either active or inactive (+1 or 0). The activity pattern at time *t* is described by 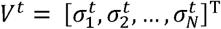 where *N* (=14) is the number of the ROIs. The *k* th brain activity pattern is described by *V*_*k*_ (*k* = 1, 2, …, 2^*N*^). Specifically, when the empirical activation of ROI *i*, ⟨*σ*_*i*_⟩, and the empirical pairwise activation of ROIs *i* and *j*, ⟨*σ*_*i*_*σ*_*j*_ ⟩, are estimated from the data, the probability distribution of the *k* th brain activity pattern with the largest entropy is the Boltzmann distribution ^55^. Here, ⟨*σ* _*i*_⟩ is equal to 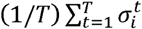 and ⟨*σ*_*i*_*σ*_*j*_⟩ is equal to 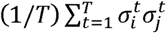, where *T* is the number of volume. That is, 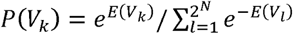, where *E*(*V*_*k*_) is the energy of activity pattern *V*_*k*_ and is given by 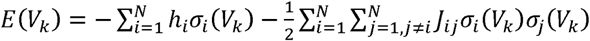. Here, *σ*_*j*_ (*V*_*k*_) represents the binarized activity (+1 or 0) at region *i* under activity pattern *V*_*k*_. Technically, *h*_*i*_ and *J*_*ij*_ in the Boltzmann distribution were adjusted until the model-based mean ROI activity 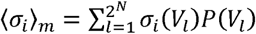 and model-based mean pairwise interaction 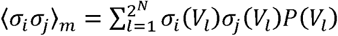 were approximately equal to the empirically obtained ⟨*σ*_*i*_⟩ and ⟨*σ*_*i*_*σ*_*j*_⟩.

#### Energy landscape analysis

We calculated the energy landscape as done in the previous studies^29,30,32,33^. The energy landscape is defined as a network of brain activity patterns *V*_*k*_ with the corresponding energy *E*(*V*_*k*_). Two activity patterns are regarded as adjacent in the network if and only if they take the opposite binary activity at just one brain region. We first exhaustively searched for local energy minimums, whose energy values are smaller than those of all the *N* adjacent patterns. We then summarized the all brain activity patterns into local minimum brain states. We first selected a starting brain activity pattern *i* among the 2^*N*^ brain activity patterns. Then, if any of its neighbor patterns has a smaller value of energy than pattern *i*, we moved to the neighbor pattern with the smallest energy value. Otherwise, we did not move, which implied that pattern *i* was a local minimum. We repeated this procedure until arrived at a local minimum. The starting pattern *i* was regarded to belong to the local minimum that was finally reached. We estimated the corresponding local minimum for all brain activity patterns.

#### Brain behavior relationship analysis

By regarding local minimum brain activity patterns as brain states, all participants have a brain state transition and behavioral time series (Fig. 1b). Thus, we can calculate behavioral performance during each brain state. We shifted the time labels of the brain state time series backwards by 5 s to account for the hemodynamic response. In addition to RT and VTC, we calculated performance accuracy (d prime). d prime was calculated as z(hit rate) - z(false alarm rate) during each brain state. Here z is normal probability density function in SciPy^56^.

### Long inter stimulus interval (ISI) gradCPT dataset (Dataset 2)

#### Participants

29 participants (13 males, ages 21–36 years, mean age = 26.4 years) performed the long inter-stimulus interval (ISI) gradCPT (1300 ms vs. 800 ms per image) during fMRI. Subjects completed the following sequence of runs with short breaks separating each (lasting a total of 1.5–2 h): one multi-echo T1-weighted run, one resting-state fMRI run, one long ISI gradCPT run followed by four long ISI gradCPT runs with intermittent thought-probes. The data used in this study and portions of the methods have been published^22^, but the current analyses and results reported have not been published elsewhere. Subjects were screened by phone and at an initial visit before the day of neuroimaging, where subjects were also trained on performing the long ISI gradCPT. Exclusion criteria were as follows: current mood, psychotic, anxiety (excluding simple phobias) or attention-deficit/hyperactivity disorder, current use of psychotropic medication, full-scale IQ less than 80, neurological disorders, sensorimotor handicaps, current alcohol or substance abuse/dependence, and claustrophobia. Furthermore, 19 ADHD patients (8 males, ages 18–34 years, mean age = 24 years) also performed the long ISI gradCPT during fMRI data collection **(Dataset 4)**. We used one long ISI gradCPT run in conjunction with four long ISI gradCPT runs with thought-probes for replication analysis but used only the four long ISI gradCPT runs with thought-probes for investigating mind wandering effect.

#### Task Paradigm and Presentation

In four long ISI gradCPT runs with thought-probes, participants performed the gradCPT, modified here to include thought-probes. The following script was used during training on a computer (outside the scanner) to instruct participants in how to respond to the thought-probes: Thought-probes appeared pseudo-randomly every 44–60 s (three possible block durations of 44, 52, and 60 s). Rather than gradually transitioning into another scene image, the last scene before the thought-probe faded into a scrambled image (to give subjects a similar amount of time to respond as in other trials). Upon the thought-probe, a question was displayed: “To what degree was your focus just on the task or on something else?” A continuous scale appeared below the question text with far-right and far-left anchors of only task and only else, respectively. Subjects pressed buttons with their middle and ring fingers to move the scale left and right, respectively, and with their thumb to enter their response. Responses were recorded on a graded scale of integers (not visible to the subjects) ranging from 0 (only task) to 100 (only else). A second self-paced question screen about meta-awareness of task-related focus (“To what degree were you aware of where your focus was?”) appeared after the thought-probe, but responses for this second question were not included in the present analyses. The gradCPT immediately resumed after subjects entered their question responses (except for the last thought-probe in the run). Scanning was manually stopped after each gradCPT thought-probe run.

#### MRI Acquisition

Functional and anatomical MRIs were acquired on the 3T Siemens CONNECTOM scanner with a custom-made 64-channel phased array head coil, housed at the Athinoula A. Martinos Center for Biomedical Imaging. The T2*-weighted whole-brain fMRI runs were performed with multiband, echo-planar imaging (simultaneous multislice factor of 4) and the following parameters: repetition time (TR), 1.08 s; echo time (TE), 30 ms; flip angle, 60°; field of view (FoV), 110 mm2; 68 transverse slices; 2 mm isotropic voxels. The T1-weighted scan parameters were as follows: TR, 2,530 ms; TE, 1.15 ms; inversion time (TI), 1,100 ms; flip angle, 7°; FoV, 256 mm2; 1 mm isotropic voxels.

### gradCPT with reward data set (Dataset 3)

#### Participants

Sixteen participants (10 males, ages = 19–29, mean age = 22 years) completed 3–5 8-min runs of the gradCPT (13 participants completed five runs, 2 completed four runs, and 1 completed three runs) during fMRI. The data used in this study and portions of the methods have been published^34^, but the current analyses and results reported have not been published elsewhere. Fourteen participants were right-handed and all were considered healthy, had normal or corrected-to-normal vision, and no reported history of major illness, head trauma, or neurological/psychiatric disorders. All were screened to confirm no metallic implants or history of claustrophobia. Drug/medication use was not explicitly assessed. The study protocol was approved by the VA Boston Healthcare System Institutional Review Board, and all participants gave written informed consent

#### Task Paradigm and Presentation

In the gradCPT with reward data set, each 8-min task run was divided into alternating 1-min rewarded and unrewarded blocks, which were differentiated by a continuous color border (green for rewarded; blue for unrewarded). To have the background colors be more intuitive and avoid confusion, “green” was chosen for rewarded blocks in all participants rather than counterbalancing green and blue colors. This yielded 4 min of each block-type per run. Similar to our previous study^35^, participants earned $0.01 for correctly pressing to city scenes and $0.10 for correctly withholding a response to mountain scenes during rewarded blocks. However, if a participant failed to press to a city scene, they would lose $0.01, and if a participant incorrectly pressed to a mountain scene they would lose $0.10. During the unrewarded blocks, no money could be gained or lost. These identical reward contingencies were shown to produce reliable improvements in accuracy and RT variability in our recent study with 54 participants^35^, and in previously published work with this data^34^, thus a priori, we were certain the payoff matrix successfully modulated sustained attention performance.

#### MRI Acquisition

Scanning was performed on a 3T Siemens MAGNETOM Trio system equipped with a 32-channel head coil at the VA Boston Neuroimaging Research Center for Veterans (NeRVe). Each gradCPT functional run included 248 whole-brain volumes acquired using an echo-planar imaging sequence with the following parameters: TR = 2000 ms, TE = 30 ms, flip angle = 90°, acquisition matrix = 64 × 64, in-plane resolution = 3.0 × 3.0 mm2, 33 oblique slices aligned to the anterior and posterior commissures, slice thickness = 3mm with a 0.75mm gap. MP-RAGE sequence parameters were as follows: TE = 3.32 ms, TR = 2530 ms, flip angle = 7°, acquisition matrix = 256 × 256, in-plane resolution = 1.0mm2, 176 sagittal slices, slice thickness = 1.0 mm.

## Code availability

All codes used in the analyses are available from the authors on request.

## Acknowledgements

This research was supported by the TOYOBO Biotechnology Foundation to AY and a Merit Review Award from the Department of Veterans Affairs Clinical Sciences Research and Development (I01CX001653) to ME.

## Author contributions

M.E., E.V. and A.K. designed the study and recruited participants for the study, collected their clinical and imaging data. A.Y. performed data preprocessing and analysis; A.Y., D.R., and M.E. primarily wrote the manuscript.

