## Supplemental Information for "Two dominant brain states reflect optimal and suboptimal attention"

**Supplementary information**

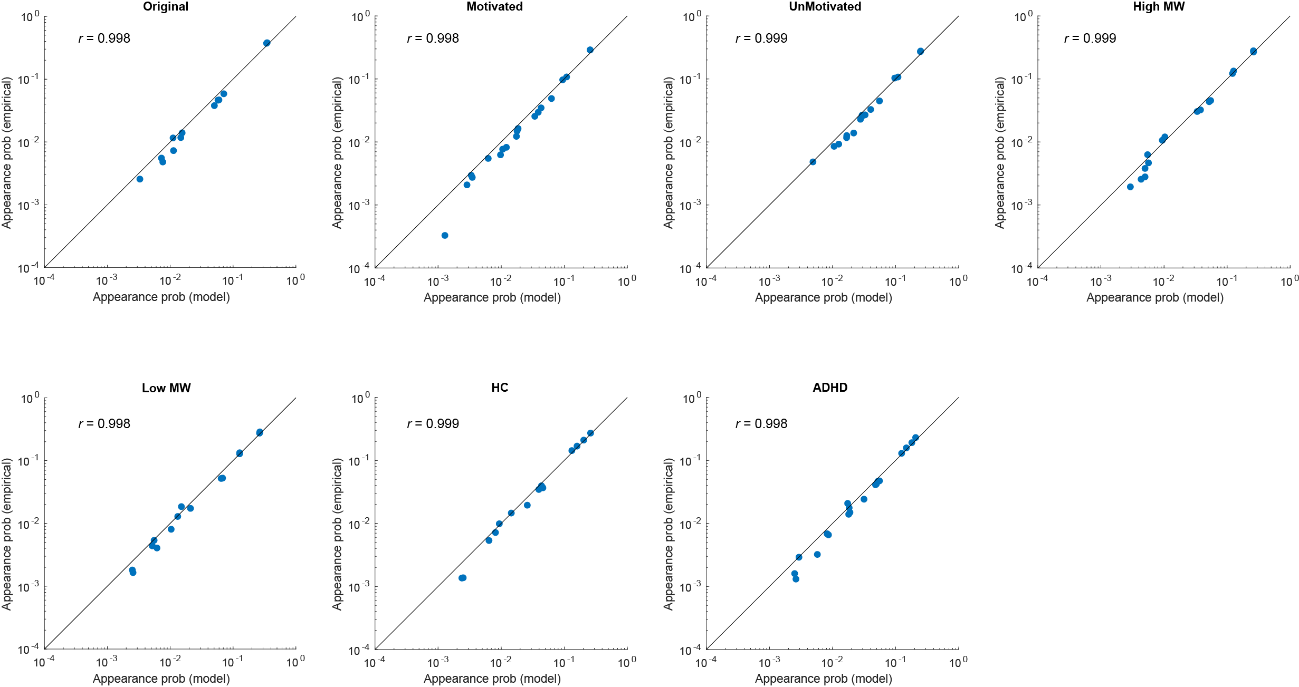

**Supplementary Figure 1. Accuracies of model fitting.** Comparison of the appearance probability of the local minimums between model and empirical for all analysis. The appearance probability of the local minimums in the empirical data was accurately reproduced by the pairwise maximum entropy model with high correlation between empirical data and model prediction (*r* > 0.99). Each scatter indicates each local minimum brain state. MW: mind wandering, AHDH: attention-deficit hyperactivity disorder.

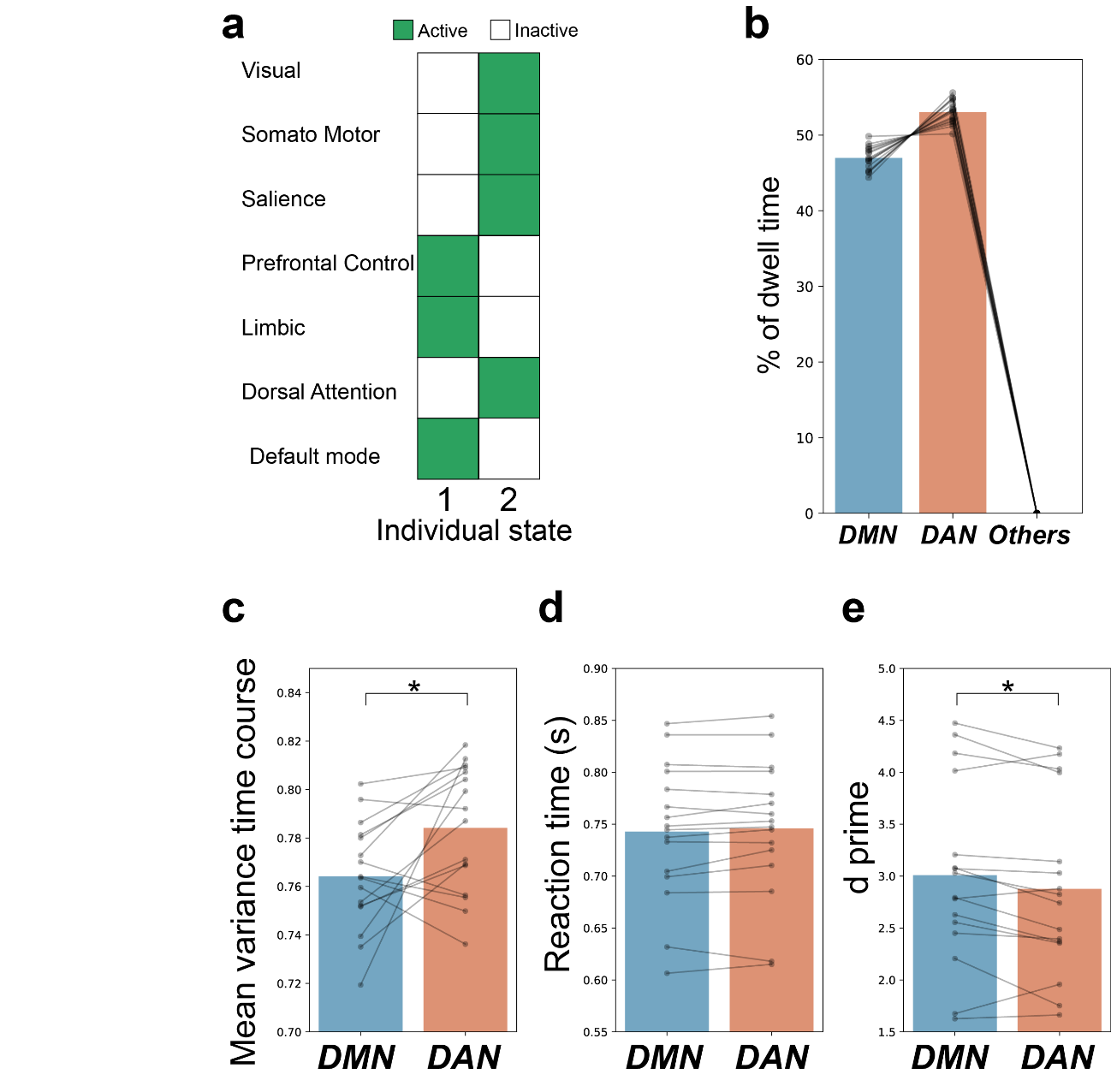

**Supplementary Figure 2.** **Replication using Schaefer et al. (REF 40) 7 regions of interest.** (a) Stable brain states during gradual onset continuous performance task (gradCPT). Individual state is represented by an activity pattern in which each brain region is active (green) or inactive (white) state. (b) Percentage of dwell time during gradCPT. Blue bars show the state defined as DMN-state, red bars show the state defined as DAN-state, green bar show the state defined as Others. (c) Mean variance time course during each state. (d) Reaction time during each state. (e) d prime during each state. Each scatter shows each participant and line connected the same participant. DMN: default mode network; DAN: dorsal attention network. **P* < 0.05 (Wilcoxon signed-rank test. Variance time course: *W_15_* = 22, *P* < 0.017; Reaction time: *W_15_* = 40, *P* > 0.14; d prime: *W_15_* = 25, *P* < 0.026, two-sided without multiple comparisons).

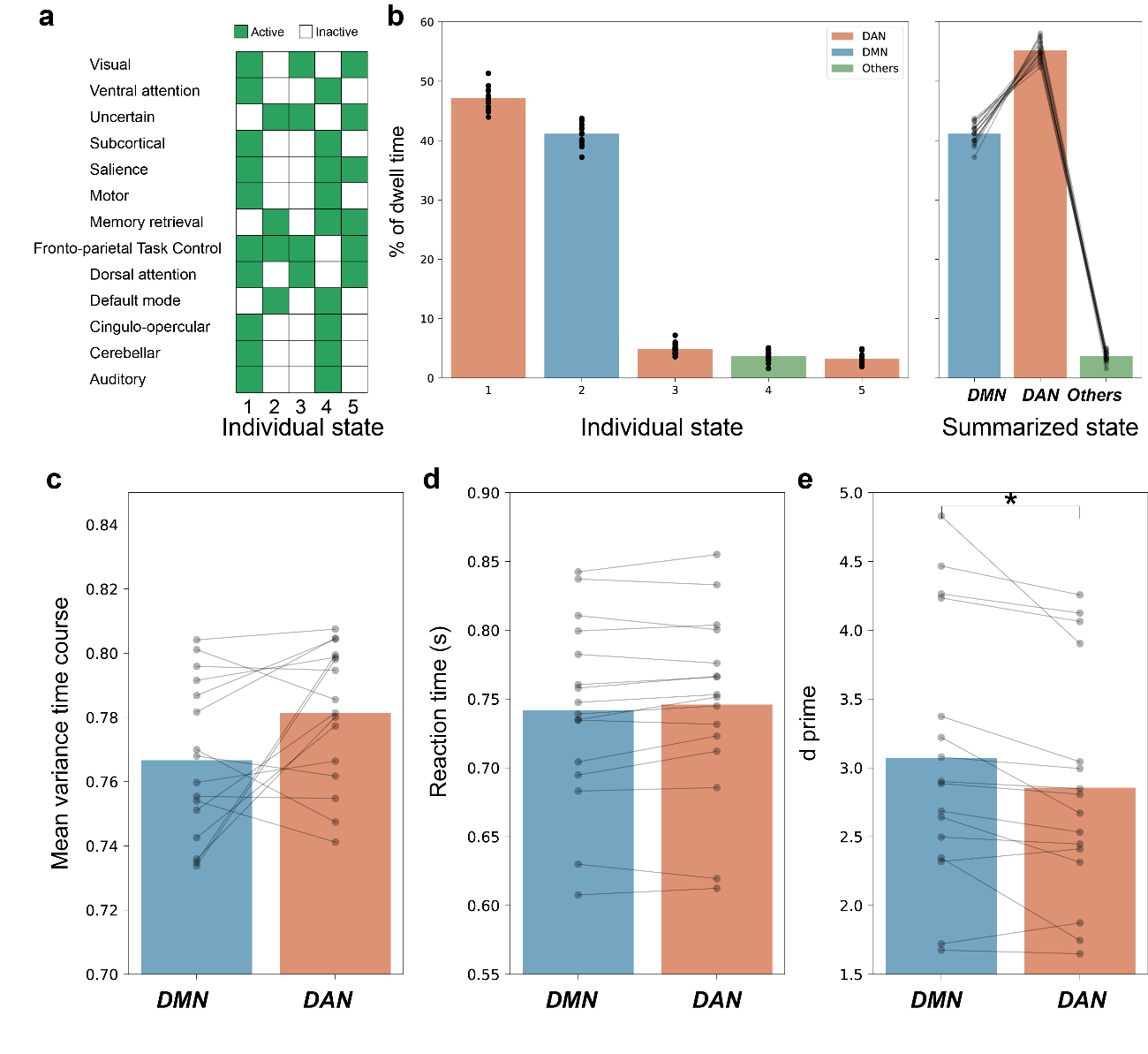
**Supplementary Figure 3. Replication using the Power’s 13 regions of interest (REF 41).** (a) Stable brain states during gradual onset continuous performance task (gradCPT). Individual state is represented by an activity pattern in which each brain region is active (green) or inactive (white) state. (b) Percentage of dwell time during gradCPT. (c) Mean variance time course during each state. Blue bars show the state defined as DMN-state, red bars show the state defined as DAN-state, green bar show the state defined as Others. (d) Reaction time during each state. (e) d prime during each state. Each scatter shows each participant and line connected the same participant. DMN: default mode network; DAN: dorsal attention network. **P* < 0.05 (Wilcoxon signed-rank test. Variance time course: *W_15_* = 32, *P* > 0.062; Reaction time: *W_15_* = 37, *P* > 0.10; d prime: *W_15_* = 14, *P* < 0.0053, two-sided without multiple comparisons).

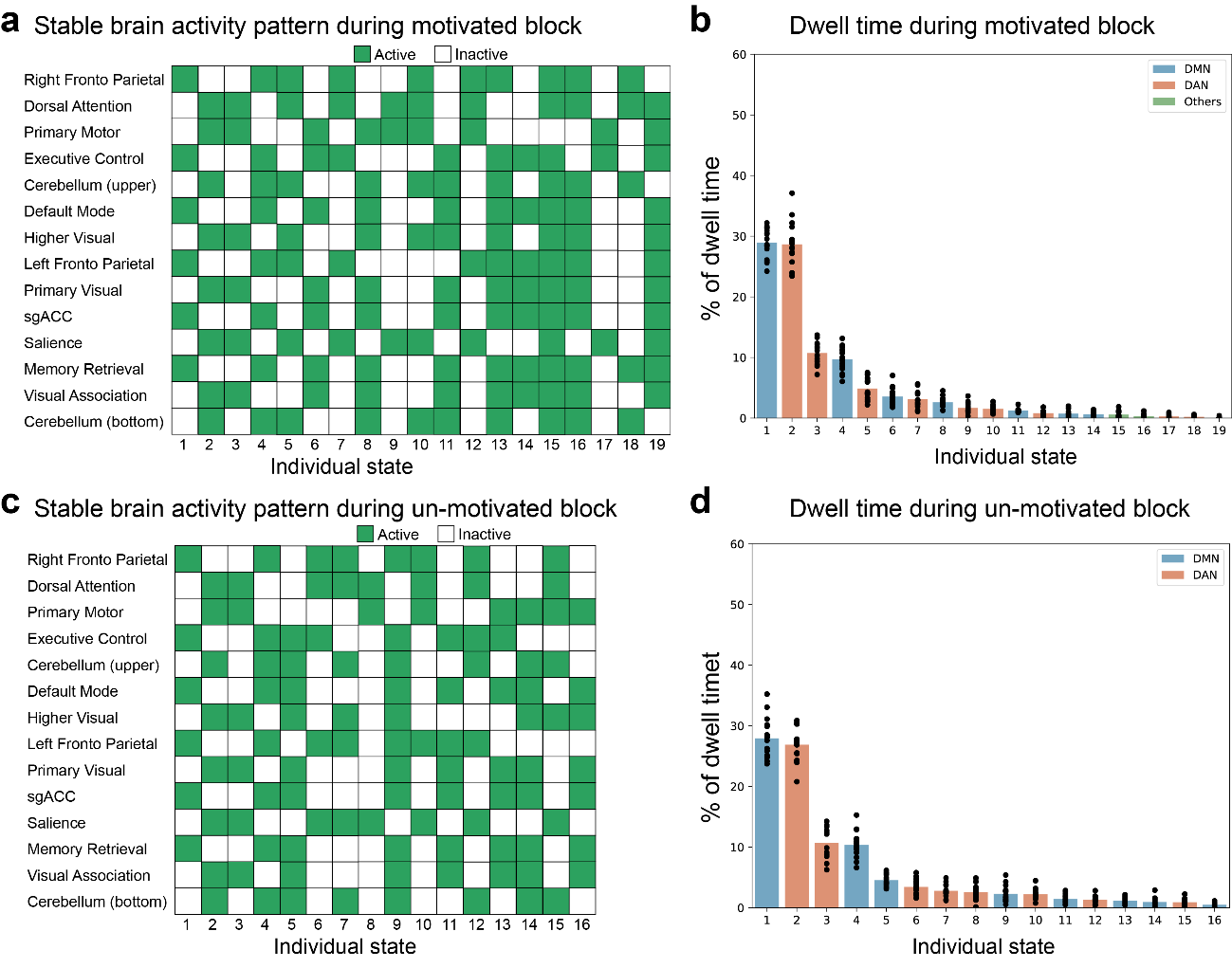

**Supplementary Figure 4.** **Energetically stable brain states and dwell time during rewarded gradCPT.** (a) Stable brain states during motivated (rewarded) blocks in gradual onset continuous performance task (gradCPT) with reward. Individual state is represented by an activity pattern in which each brain region is active (green) or inactive (white) state. (b) Left graph shows percentage of dwell time during gradCPT in each individual state. Blue bars show the state defined as DMN-state, red bars show the state defined as DAN-state, green bar show the state defined as Others. Each scatter shows each participant. (c,d) Stable brain states and percentage of dwell time during unmotivated (unrewarded) blocks. DMN: default mode network; DAN: dorsal attention network.

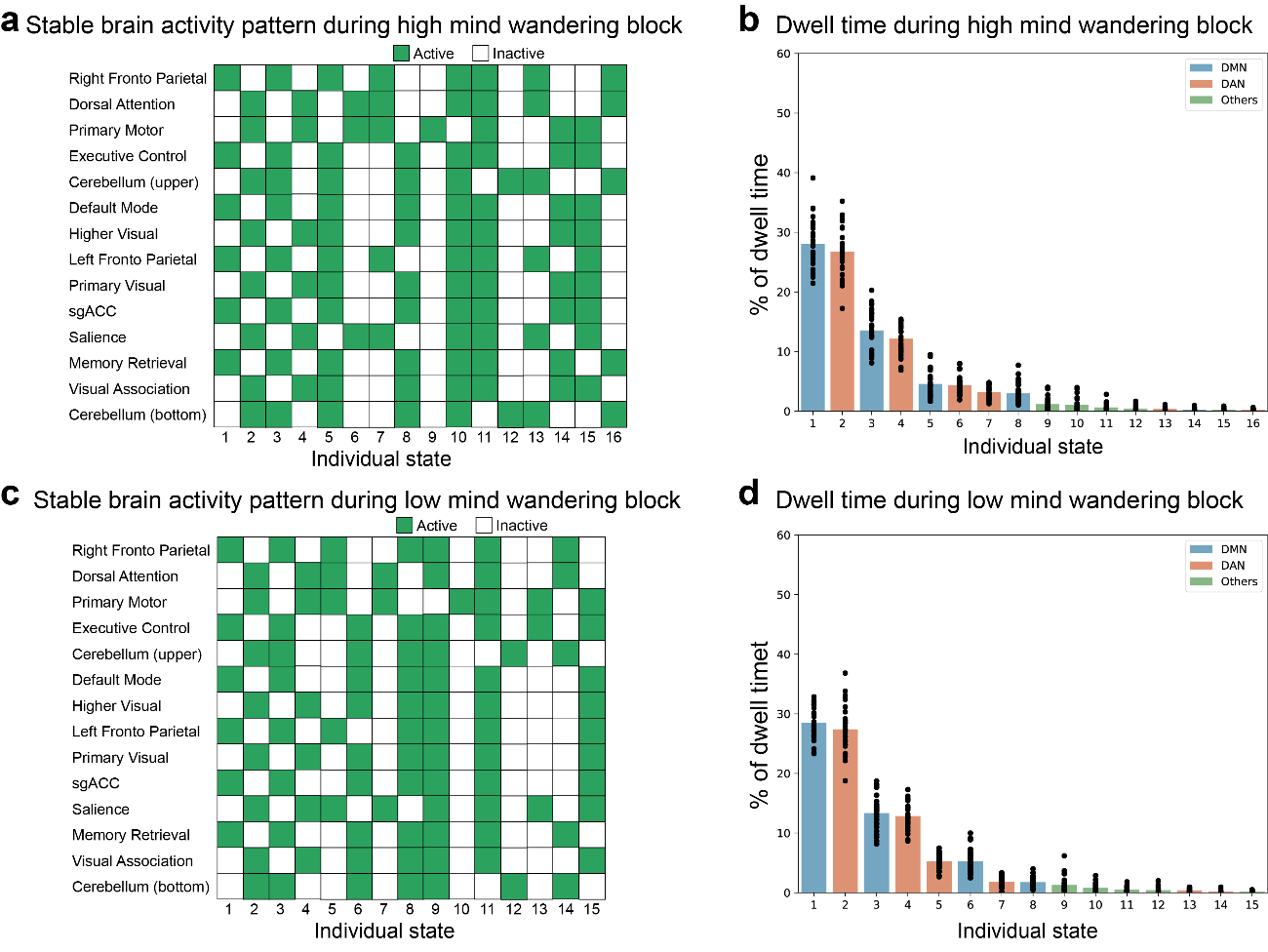

**Supplementary Figure 5. Energetically stable brain states and dwell time during gradCPT with mind wandering probes.** (a) Stable brain states during high mind wandering blocks in gradual onset continuous performance task (gradCPT) with mind wandering probes. Individual state is represented by an activity pattern in which each brain region is active (green) or inactive (white) state. (b) Left graph shows percentage of dwell time during gradCPT in each individual state. Blue bars show the state defined as DMN-state, red bars show the state defined as DAN-state, green bar show the state defined as Others. Each scatter shows each participant. (c,d) Stable brain states and percentage of dwell time during low mind wandering blocks. DMN: default mode network; DAN: dorsal attention network.

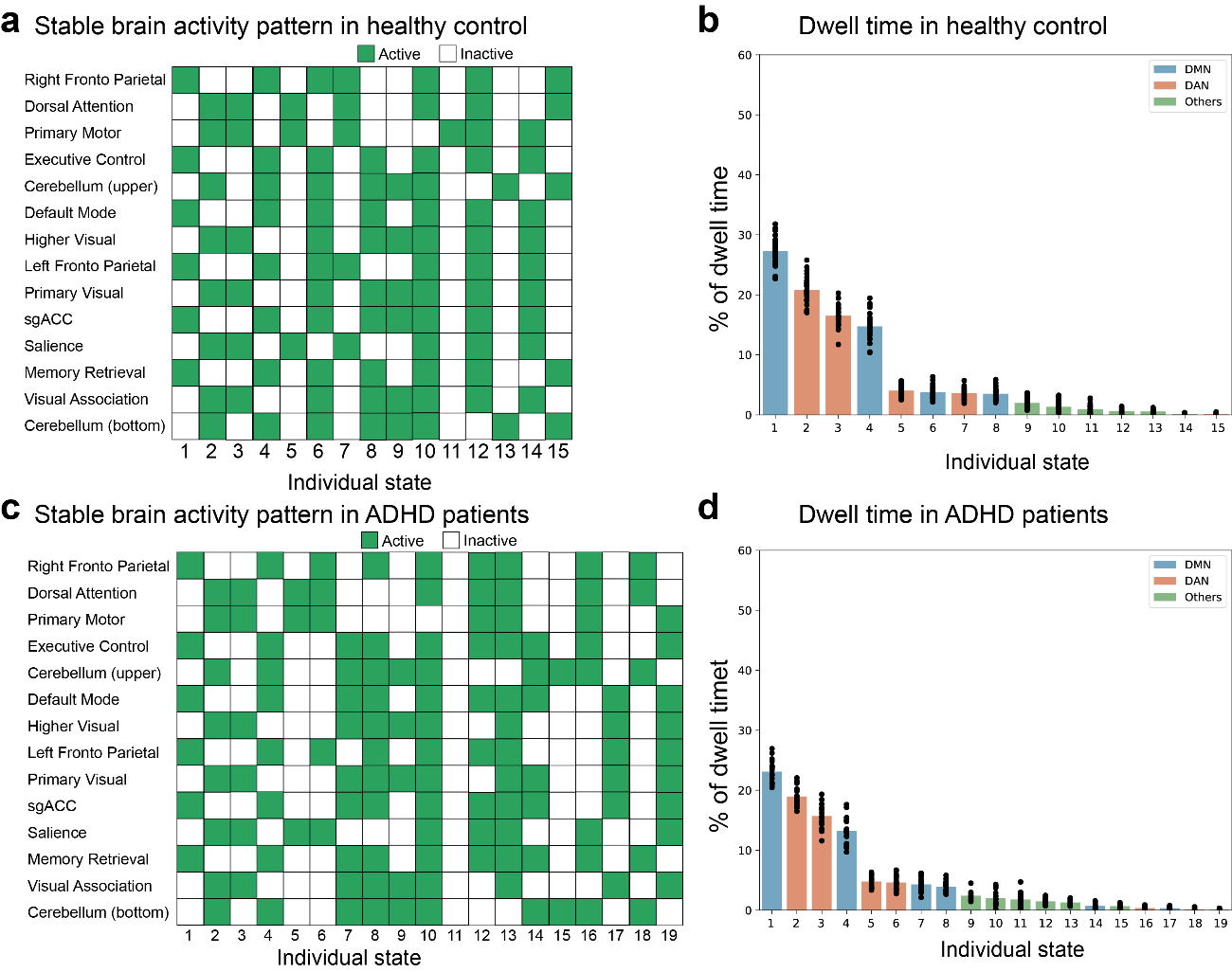

**Supplementary Figure 6. Energetically stable brain states and dwell time for healthy control and ADHD patient.** (a) Stable brain states during gradual onset continuous performance task (gradCPT) in healthy control. Individual state is represented by an activity pattern in which each brain region is active (green) or inactive (white) state. (b) Left graph shows percentage of dwell time during gradCPT in each individual state. Blue bars show the state defined as DMN-state, red bars show the state defined as DAN-state, green bar show the state defined as Others. Each scatter shows each participant. (c,d) Stable brain states and percentage of dwell time in ADHD patients. DMN: default mode network; DAN: dorsal attention network, AHDH: attention-deficit hyperactivity disorder.
